# Exome wide association study on Albuminuria identifies a novel rare variant in *CUBN* and additional genes, in 33985 Europeans with and without diabetes

**DOI:** 10.1101/355990

**Authors:** Tarunveer S. Ahluwalia, Christina-Alexendra Schulz, Johannes Waage, Tea Skaaby, Niina Sandholm, Natalie van Zuydam, Romain Charmet, Jette Bork-Jensen, Peter Almgren, Betina H. Thuesen, Mathilda Bedin, Ivans Brandslund, Cramer K. Chrisitansen, Allan Linneberg, Emma Ahlqvist, Per-Henrik Groop, Samy Hadjadj, David-Alexandre Tregouet, Marit E. Jørgensen, Niels Grarup, Matias Simons, Leif Groop, Marju-Orho Melander, Mark McCarthy, Olle Melander, Peter Rossing, Tuomas O. Kilpelainen, Torben Hansen

**Author notes:** Equal contributions (Tuomas O. Kilpelainen, Torben Hansen). **Corresponding author:** Tarunveer Singh Ahluwalia, PhD, Senior Researcher, Steno Diabetes Center Copenhagen, Herlev and Gentofte Hospital, The Capital Region, Copenhagen, Denmark.

## Abstract

Identifying rare coding variants associated with albuminuria may open new avenues for preventing chronic kidney disease (CKD) and end-stage renal disease which are highly prevalent in patients with diabetes. Efforts to identify genetic susceptibility variants for albuminuria have so far been limited with the majority of studies focusing on common variants.

We performed an exome-wide association study to identify coding variants in a two phase (discovery and replication) approach, totaling to 33,985 individuals of European ancestry (15,872 with and 18,113 without diabetes) and further testing in Greenlanders (n = 2,605). We identify a rare (MAF: 0.8%) missense (A1690V) variant in *CUBN* (rs141640975, β=0.27, p=1.3 × 10^−11^) associated with albuminuria as a continuous measure in the combined European meta-analyses. Presence of each rare allele of the variant was associated with a 6.4% increase in albuminuria. The rare *CUBN* variant had 3 times stronger effect in individuals with diabetes compared to those without *(pinteraction:* 5.4 × 10^−4^, β_DM_: 0.69, β_*nonDM:*_ 0.20) in the discovery meta-analyses. Gene-aggregate tests based on rare and common variants identify three additional genes associated with albuminuria *(HES1, CDC73*, and *GRM5)* after multiple testing correction (*P*__bonferroni_<2.7 × 10^−6^).

The current study identifies a rare coding variant in the *CUBN* locus and other potential genes associated with albuminuria in individuals with and without diabetes. These genes have been implicated in renal and cardiovascular dysfunction. These findings provide new insights into the genetic architecture of albuminuria and highlight novel target genes and pathways for prevention of diabetes-related kidney disease.

**Significance statement:** Increased albuminuria is a key manifestation of major health burdens, including chronic kidney disease and/or cardiovascular disease. Although being partially heritable, there is a lack of knowledge on rare genetic variants that contribute to albuminuria. The current study describes the discovery and validation, of a new rare gene mutation (~1%) in the *CUBN* gene which associates with increased albuminuria. Its effect multiplies 3 folds among diabetes cases compared to non diabetic individuals. The study further uncovers 3 additional genes modulating albuminuria levels in humans. Thus the current study findings provide new insights into the genetic architecture of albuminuria and highlight novel genes/pathways for prevention of diabetes related kidney disease.

## Introduction

Albuminuria is a manifestation of chronic kidney disease (CKD), a major health burden worldwide with a current prevalence of 14.8% in the United States^1^. In individuals with CKD, changes in albuminuria are strongly associated with the risk of end-stage renal disease and death^2^. Albuminuria has also been associated with increased risk of cardiovascular disease and mortality among individuals with and without diabetes^3–5^.

Family-studies suggest that genetic factors explain 16% to 49% of albuminuria ^6^. While several genome wide association studies (GWAS) of albuminuria have been performed to date, most have focused on identifying common genetic variants (minor allele frequency (MAF) ≥ 5%) for albuminuria ^6–8^. In a recent study, we identified rare coding variants for kidney function (estimated glomerular Alteration rate, eGFR) and development using exome-wide association analyses ^9^ Here, we use a similar approach to identify rare (MAF <1%) or low frequency (MAF 1-5%) coding variants for albuminuria in 33,985 individuals of European ancestry with (n=15,872) and without (n=18,111) diabetes.

## Materials and Methods

### Study populations

The present study comprises a two-stage design: discovery and replication. The discovery set includes five cohorts from Denmark (Inter99, Health-2006, Health-2008, Vejle Biobank, and Addition-DK) with a total of 13,226 participants (3,896 with and 9,330 without diabetes), described previously ^10, 11^**(Supp. Text 1.1)**.

The replication set includes multiple studies of European (EUR) descent (*n*_EUR_: 20,759; 11,976 with and 8,783 without diabetes) including the IMI-SUMMIT consortia (Europe-UK based consortia on diabetes cases)^12^, and Greenlandic (GL) Inuit populations (*n*_GL_: 2,605) which are all described in detail in **Supp. Text 1.2**. The overall study design involving the single SNP association testing and Gene based aggregate testing has been presented in form of process flow diagrams **(Figure 1A and 1B**).

**Figure 1.**
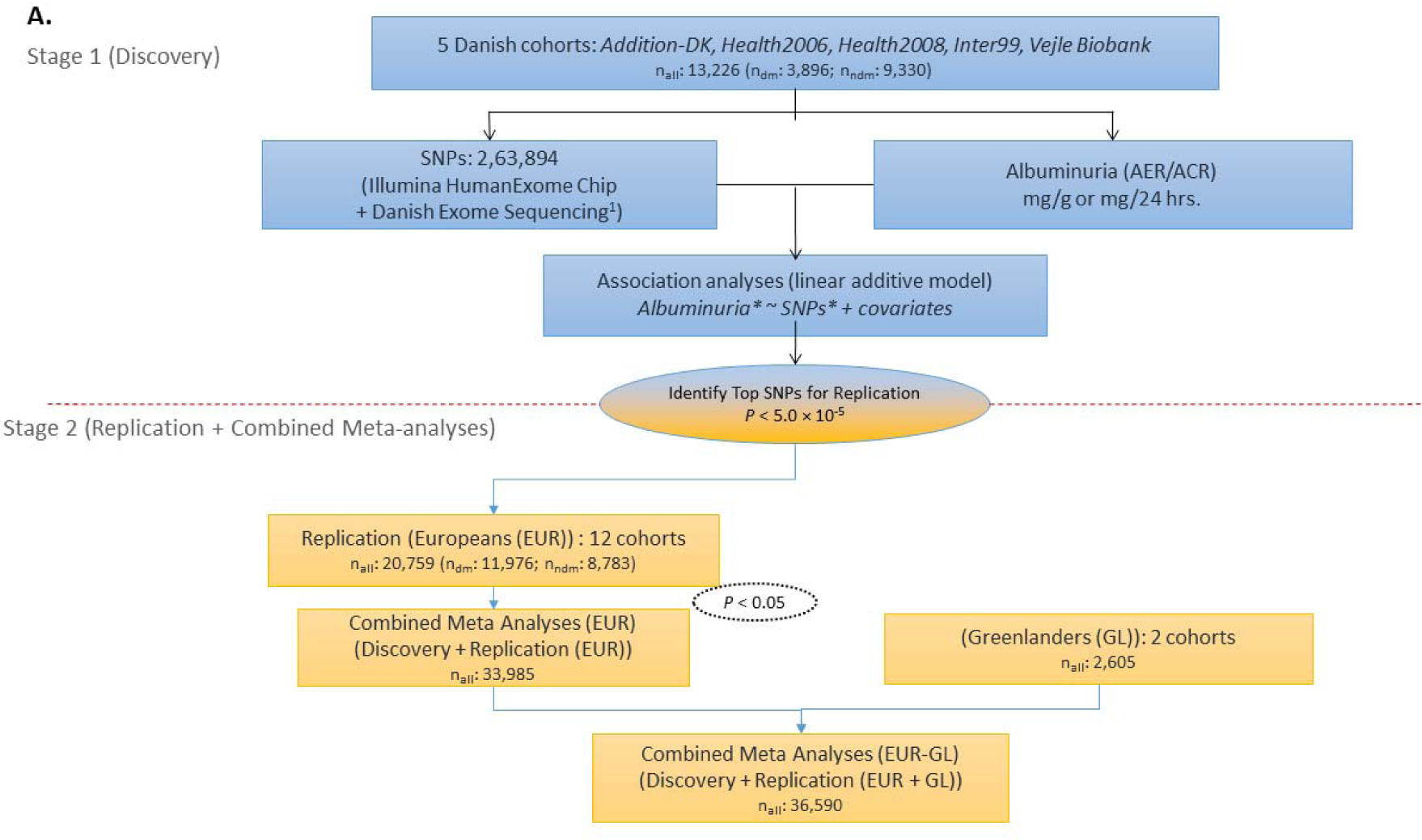
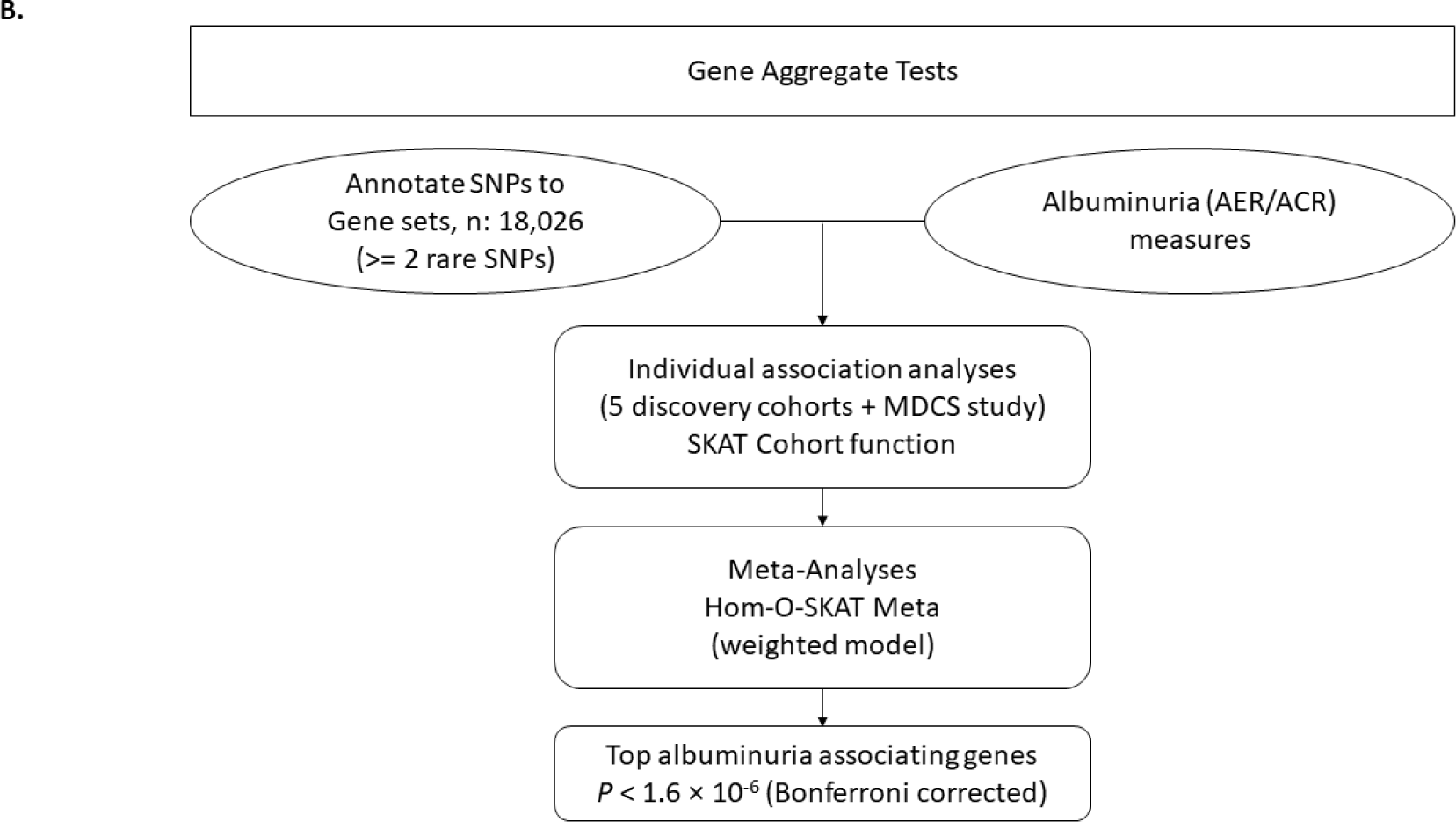
Study Design Overview: A. Exome wide association study (Ex-WAS) on Albuminuria: A multi stage study on Europeans and Greenlanders

The present study was conducted in accordance with the Helsinki declaration and all the participating studies were approved by their respective Data Protection Boards and by the regional scientific ethics committees. All participants signed a written consent.

### Phenotype measurements

Albuminuria was diagnosed from a 24-hour urine collection (mg/24 hours), also called the urinary albumin excretion rate (AER) or, from spot urine samples measuring urinary albumin and creatinine concentrations and calculating their ratio (ACR in mg/g). The methods used to measure AER/ACR by participating cohorts have been described in the **Supp. table S1.1** with phenotype measures summary in **Table 1**.

**Table 1.** Clinical characteristics of the subjects for individual cohorts: Pooled and stratified on diabetes status: Discovery and Replication stages.

| Study Name | Sample Set Type | N | Women, % | Age *, years | ACR<sup>a,*</sup> (mg/g) or AER<sup>b,*</sup> (mg/24Hrs) | Diabetes, % | Diabetes, Type | Diabetes duration, years<sup>*</sup> |
| --- | --- | --- | --- | --- | --- | --- | --- | --- |
| Stage 1 (Discovery) |  |  |  |  |  |  |  |  |
| Addition-DK | All | 2013 | 47.3 | 59.6 (7.0) | 5.3 (1.8-14.1)<sup>a</sup> | 81.6 | Type 2 | - |
|  | DM | 1,643 | 43.9 | 60.1 (6.8) | 5.3 (1.8-15.0)<sup>a</sup> |  |  | - |
|  | Non-DM | 370 | 62.4 | 57.6 (7.4) | 4.4 (1.8-12.3)<sup>a</sup> |  |  | - |
| Health2006 | All | 2,658 | 51.3 | 49.6 (12.8) | 5.0 (4.0-8.0)<sup>a</sup> | 0 | - | - |
| Health2010 | All | 642 | 55.6 | 46.6 (8.2) | 4.0 (3.0-5.0)<sup>a</sup> | 0 | - | - |
| Inter99 | All | 5,971 | 50.9 | 46.1 (7.9) | 3.0 (2.0-5.0)<sup>a</sup> | 5.2 | Type 2 | - |
|  | DM | 311 | 37.6 | 51.0 (7.1) | 4.0 (3.0-10.0)<sup>a</sup> |  |  | - |
|  | Non-DM | 5,660 | 51.6 | 45.9 (7.8) | 3.0 (2.0-5.0)<sup>a</sup> |  |  | - |
| Vejle Biobank | DM | 1,942 | 38.1 | 63.4 (8.7) | 8.5 (4.8-16.2)<sup>b</sup> | 100 | Type 2 | - |
| Stage 2 (Replication 1): EUROPEANS |  |  |  |  |  |  |  |  |
| DanFUNd | All | 7,364 | 47.3 | 52.1 (13.1) | 5.0 (5.0-5.0)<sup>a</sup> | 6.3 | Type 2 | - |
|  | DM | 449 | 43.9 | 60.4 (8.8) | 5.0 (5.0-13.0)<sup>a</sup> |  |  | - |
|  | Non-DM | 6,915 | 62.4 | 51.4 (13.3) | 5.0 (5.0-5.0)<sup>a</sup> |  |  | - |
| Genesis/Genediab | DM | 1,249 | 52.2 | 42.2 (11.9) | 10 (4.5-120)<sup>a</sup> | 100 | Type 1 | 25.6 (10.2) |
| MDCS | All | 2,641 | 56.5 | 73.0 (5.6) | 5.3 (3.5-10.6)<sup>a</sup> | 21 | Type 2 | - |
|  | DM | 547 | 48.3 | 73.6 (5.4) | 7.1 (3.5-19.5)<sup>a</sup> |  |  | - |
|  | Non-DM | 2,094 | 58.6 | 72.9 (5.6) | 5.3 (3.5-9.7)<sup>a</sup> |  |  | - |
| SUMMIT Consortium (Diabetes cohorts) |  |  |  |  |  |  |  |  |
| Benedict (Phase A and B) | DM | 324 | 32.7 | 68.1 (7.7) | 17.3 (4.6-55.6)<sup>b</sup> | 100 | Type 2 | 17.2 (7.1) |
| Cambridge | DM | 245 | 46.9 | 23.1 | 6.9 (4.6- | 100 | Type 1 | 15.7 (6.5) |
|  |  |  |  | (9.9) | 15.1)<sup>a</sup> |  |  |  |
| **Eurodiab** | DM | 680 | 50.0 | 42.5<br>(9.9) | 23.9 (22.6-25.3)<sup>b</sup> | 100 | Type 1 | 24.7 (8.4) |
| **FinnDiane** | DM | 2,840 | 50.6 | 43.9<br>(11.7) | 11.0 (5.0-89.6)<sup>b</sup> | 100 | Type 1 | 29.1<br>(10.0) |
| **GoDarts 1** | DM | 3,530 | 46.0 | 67.0<br>(0.7) | 408.1<br>(342.9-466.1)<sup>a</sup> | 100 | Type 1 | 7.5 (6.0) |
| **GoDarts 2** | DM | 2,805 | 43.0 | 66.8<br>(11.8) | 447.4<br>(360.8-525.3)<sup>a</sup> | 100 | Type 2 | 7.3 (6.2) |
| **SDR (Type1)** | DM | 598 | 43.0 | 48.6<br>(13.7) | 8.6 (4.3-50.4)<sup>b</sup> | 100 | Type 1 | 31.9<br>(13.3) |
| **SDR (Type2)** | DM | 1,426 | 41.3 | 65.6<br>(10.7) | 18.7 (7.2-100.8)<sup>b</sup> | 100 | Type 2 | 14.3 (7.6) |
| **Steno Type 2**<br>**Diabetes** | DM | 295 | 39.3 | 61.5<br>(8.1) | 50.5 (13.9-893.0)<sup>b</sup> | 100 | Type 2 | 15.1 (6.8) |
| **Stage 2 (Replication 2): GREENLANDERS** |  |  |  |  |  |  |  |  |
| **Greenlanders**<br>**(IHIT + B99)** | All | 2,605<br>(IHIT=2,519;<br>B99=86) | 53.7 | 44.1<br>(14.5) | 8.0 (6.0-14.0)<sup>a</sup> | 8.1 | - | - |

\*Age and diabetes duration are in mean ± standard deviation while ACR (mg/g)/AER (mg/24 hrs.) are represented as Median (IQR: Inter quartile range). For some studies AER was converted from $\mu$ g/min to mg/24 hours by a multiplication factor of 1.44 ( $\mu$ g/minute × 1.44= mg/24 hours or mg/g). Sample Set Type is represented as All: All the individuals in the cohort. All= DM + Non-DM. These have been further stratified based on presence (DM) or absence of diabetes (Non-DM). ACR: Urinary albumin-creatinine ratio, AER: albumin excretion rate; DM: Diabetes Mellitus. The number of individuals in the phenotyping summary in this table may not match the association summary numbers in actual analyses due to missing genetic information for some individuals.

### Genotyping and SNP quality control

Genotyping of the discovery phase studies was performed on the Illumina HumanExome BeadChip 12V. 1.0 containing 2,44,883 single nucleotide polymorphisms (SNPs) plus additional ~17,000 custom-typed SNPs from the Danish Exome Sequencing Project as described previously ^13^. The total number of SNPs was 263,894 of which most were exome based (non-synonymous/coding) gene variants (~90%). Thus, we refer to the present association study as an exome-wide association study (Ex-WAS).

Genotyping methods and quality control (QC) based inclusions and exclusions in the discovery and replication (20,759 EUR and 2,605 GL) cohorts have been described in **Supp. table S1.2.**

Albeit a total of 54,606 SNPs remained for a total of 13,226 individuals with complete phenotype and genotype data in the discovery set.

## Statistical analyses

### Discovery Phase meta Ex-WAS

The discovery phase Ex-WAS for albuminuria was first performed in each of the five participating studies individually using additive linear regression model adjusting for gender, age, and population sub-structure (first ten principal components), and systolic blood pressure (SBP) wherever available. Albuminuria measurements were natural log transformed to correct for non-normalised data.

The study-specific results were meta-analysed (meta Ex-WAS) using inverse variance-weighted fixed effects meta-analyses where weights are proportional to the squared standard errors of the effect estimates. Genomic inflation factor (λ) was at acceptable levels both in the individual study ^Ex-WAS^ (λ_inter99_ = ^1.01^, λ_health2006_ = ^1.0, λhealth2008^ = ^1.0^, λ_vejle_ = ^0.99,^ λ_addition_dk_ = ^1.0^1) 3^nd^ in the combined discovery meta Ex-WAS λ_discovery_=10). A chi-square test for heterogeneity was implemented to estimate the heterogeneity in effect sizes across the participating studies. The METAL software (http://csg.sph.umich.edu/abecasis/metal/) was used for meta-analyses and the R meta package (https-//github.com/guido-s/meta) for constructing meta forrest plots (discovery set). Ancestry specific linkage disequilibrium (LD) between variants was extracted using the NIH based LDlink database (https://analysistools.nci.nih.gov/LDlink/) or through the SNAP (**SN**P **A**nnotation and **P**roxy Search) (http://archive.broadinstitute.org/mpg/snap/index.php) database.

### Replication Phase meta analyses

SNPs with *p* < 5.0 × 10^−5^ in the discovery meta-analyses were tested for replication in European *(n*_*EUR*_*: 20,759)* and Greenlandic *(n*_*GL*_*: 2,605)* study cohorts independently using the linear regression models (adjusting for study specific covariates). SNPs with imputation quality r^2^ <0.3 were not used for replication. Following this an inverse variance or *p* value based fixed effects meta-analyses was run (wherever required). Replication meta-analyses with *p*_*replication*_ <0.05, were considered significant.

### Combined (EUREUR-GL) Meta analyses

The combined meta-analyses was firstly performed with all individuals of European ancestry (Combined EUR) followed by pooling of Greenlandic data (Combined EUR-GL).

Any SNP with a) *p*_replication_ < 0.05 and b) *p*_meta_EUR/EUR-GL_ < 5.0 × 10^−8^ was considered overall significant while those with 5.0 × 10^−5^ &#x003E; *p*_meta_EUR/EUR-GL_ > 5.0 × 10^−8^ were considered suggestive.

### Conditional analyses

Conditional analyses for novel SNPs identified in known loci (and/or in low LD, LD_r2_<0.01) were performed to determine if the signal was independent (in discovery set). This was performed using the following linear model: *trait ~ top identified SNP + secondary known SNP + other covariates.* If the top SNP retained the association estimates and *p* value it was considered an independent signal.

### Gene Aggregate Tests

Gene-based multi-marker association testing for rare and common (MAF > 0.0001) variants was performed using the Meta Analyses for SNP-Set (Sequence) Kernel Association Test (MetaSKAT) R package (https://cran.r-project.org/web/packages/MetaSKAT/index.html). At the study-specific level, the gene-based analyses were performed against the null model accounting for gender and ten principal components, (using the previously described SKAT-O method) ^14^ generating SKAT objects individually for each cohort with complete ExomeWAS data (discovery studies + MDCS study, *n*_*studies*_ = 6)) which were then meta-analyzed. The meta-analyses of the summary-level score statistics were run using the Hom-Meta-SKAT-O “optimal” method (optimal linear combination of burden and SKAT statistics) which assumes that different studies share the same causal variant, weighing them equally. An alternate method, Het-Meta-SKAT-O, which is based on the assumption that studies/groups may have different causal variants, was also tested. Genes with less than 2 contributing variants were filtered out. A Bonferroni-based correction was applied on 18, 026 gene sets tested (*P*_adjusted_<2.7 × 10^−6^).

### Phenome Scanning: SNP-trait associations from publicly available data (GWAS, eQTL based gene expression, and SNP-Metabolite associations)

The Pheno Scanner database (http://www.phenoscanner.medschl.cam.ac.uk/phenoscanner) comprising publicly available results for SNP-trait associations (GWAS, eQTL and metabolomics data) for the European ancestry ^15^ was used to mine known SNP-Trait associations of the current SNP findings.

### SNP functionality prediction: SIFT

We used the SIFT (Sorting Intolerant from Tolerant) database to check whether any top coding variant was associated with functional by nature (http://sift.bii.a-star.edu.sg/).

## RESULTS

### SNP-Albuminuria Association Analyses Discovery Phase

In stage 1 discovery phase meta-analyses comprising five Danish studies *(n*_*total*_. 13,226), three independent SNPs reached *p*_*discovery*_ < 5.0 × 10^−5^. This included two rare variants (MAF<0.01) in *CUBN* and *LMX1B* and a common variant in *KCNK5.* The quantile-quantile (QQ) and manhattan plot (stage 1) can be found in the supplementary material **(Supp. figures S1a and S1b**) along with the locus zoom plots for the 3 top SNP signals (**Supp. figures S2a, S2b, and S2c**).

Top signal in *CUBN* (cubilin) gene rs141640975 is a rare (MAF: 0.0083) missense (A1690V) SNP with the A allele associated with increased albuminuria (0: 0.25; *p*_*discovery*_: 1.2 × 10^−5^, **Table 2, Supp. figure S3**). The *KCNK5* (potassium two pore domain channel subfamily K member 5) common (MAF: 0.23) intronic SNP rs10947789 minor (C) allele (0: 0.05; *p*_*discovery*_: 1.6 × 10^−5^, **Table 2, Supp. figure S4**) and the *LMX1B* (LIM homeobox transcription factor 1 beta) rare (MAF: 0.0089) intronic SNP rs140177498 T allele associated with increased albuminuria (0: 0.26; *p*_*discovery*_: 8.7 × 10^−6^, **Table 2, Supp. figure S5**).

**Table 2.** Association analyses results on top SNPs from Ex-WAS discovery, replication and combined meta-analyses including Europeans (EUR) and Europeans + Greenlanders (EUR + GL), for albuminuria.

| SNP Characteristics |  |  |  |  | Discovery<br>*n* = 13, 226 |  |  |  | Replication<br>*n* = 20, 759 |  |  | COMBINED<br>(EUR)<br>*n* = 33, 985 |  | COMBINED<br>(EUR and GL)<br>*n* = 36, 590 |  |  |
| --- | --- | --- | --- | --- | --- | --- | --- | --- | --- | --- | --- | --- | --- | --- | --- | --- |
| SNP<br>(rsID)* | Gene | Chr<br>(Pos) | Anno. | EA/<br>OA | EA<br>F | BET<br>A | SE | P<sub>discov</sub><br>ery | N | BET<br>A | P<sub>replica</sub><br>tion | N | P<sub>combi</sub><br>ned | N | P<sub>me</sub><br>ta | P<sub>he</sub><br>t |
| **Novel SNP ($P_{meta} < 5.0 \times 10^{-8}$)** |  |  |  |  |  |  |  |  |  |  |  |  |  |  |  |  |
| rs141640975 | *CUBN* | 10<br>(16992011) | missense<br>(A1690V) | A/G | 0.008 | 0.266 | 0.061 | 1.2e-05 | 9,742 | 0.279 | 2.9e-07 | 22,866 | 1.3e-11 | - | - | - |
| **Suggestive SNP ($5.0 \times 10^{-5} > P_{meta} > 5.0 \times 10^{-8}$)** |  |  |  |  |  |  |  |  |  |  |  |  |  |  |  |  |
| rs10947789 | *KCNK5* | 6<br>(39174922) | Intronic | C/T | 0.23 | 0.054 | 0.012 | 1.6e-05 | 20,759 | - | 0.03 | 33,985 | 1.5e-05 | 36,590 | 9.1e-06 | 0.84 |

SNPs with $P_{meta} < 5.0 \times 10^{-8}$ are novel and $5.0 \times 10^{-8} < P_{meta} < 5.0 \times 10^{-5}$ are suggestive. \*SNPs were selected for replication based on $P_{discovery} < 5.0 \times 10^{-5}$ . Abbreviations: SNP: Single nucleotide polymorphism; Chr: Chromosome, Pos: SNP Position in base pairs from the GRCh37 assembly build in dbSNP; EA/OA: Effect allele/other allele, EAF: effect allele frequency, SE: standard error, P: P value for association, N: number of individuals in the analyses, EUR: European ancestry (discovery + replication), GL: Greenlanders, EUR and GL (Europeans and Greenlanders combined meta-analyses). Discovery set is based on n upto 13,226 individuals (3,896 with and 9,330 without diabetes). BETA values correspond natural log transformed Albuminuria levels (ACR:mg/g or AER:mg/24 hours).

### Diabetes stratified SNP-albuminuria effects

Effect of diabetes status on SNP-albuminuria association was assessed in the discovery set (3,896 diabetic (DM) and 9,330 non diabetic (NDM) individuals) after pooling individual genotype data on all participants, through an interaction model. Replication studies were not included in this analyses due to non-availability of individual level genotype data.

Significant inter-group (DM and Non-DM) differences in effect estimates were observed through the interaction model *(trait ~ SNP + DMstatus + SNP*DMstatus)* with a *P*_*interaction*_- 5.4 × 10^−4^.

Effect estimate of *CUBN*rs141640975 were more than 3 folds in the DM group (β: 0.69; *p*_*DM*_: 2.0 × 10^−5^ **, Supp. figure S3a**) in comparison to the Non-DM group (**Supp. table S1.3, Supp. figure S3b**). There was no significant interaction effect detected for the other 2 SNPs (rs 10947789 and rs140177498, *P*_interaction_>0.05). Individual SNP effects in DM stratified analyses as forrest plots for *KCNK5* and *LMX1B* (DM and NDM) are depicted in the Supplementary material (**Supp. figures S4a, S4b, S5a, and S5b**).

### Replication Phase

Replication of the three SNPs *(CUBN* rs141640975, *LMX1B* rs140177498 and *KCNK5* rs10947789) was sought in up to 20,759 individuals (n_DM_: 11,976; *n_non_-_DM_:* 8,783) with measures on albuminuria. The *CUBN* rs141640975 replicated strongly (*n*_*replication_CUBN*_: 9,742; *p*_*replication*_: 2.8 × 10^−7^), while the *KCNK5* rs10947789 replicated nominally *(n*_*repilcation_KCNK5*_: 20,757; *p*_*replication*_: 0.03). The *LMX1B* rs140177498 did not replicate *(n*_*replication_LMXiB*_ 13,233; *p*_*replication:*_ 0.43) (**Table 2)** and was not analyzed any further.

### Combined Meta-analyses (EUR and EUR-GL)

The combined meta-analyses (EUR: discovery + replication) comprised a total of 33,985 EUR individuals while the EUR-GL comprised 36,590 individuals in the pooled set (**Table 2**). Only *KCNK5* SNP was available in the Greenlandic data for replication.

The *CUBN* rs141640975 remained significant after the combined EUR meta-analyses *(p*_*meta_EUR*_. 13 × 10^−11^), whereas the *KCNK5* SNP reached suggestive significance (5.0 × 10^−5^ > *P*_meta_EUR-GL_ < 5.0 × 10^−8^) in the EUR only and EUR-GL meta-analyses (**Table 2**).

### Conditional analyses

Conditional analyses for the identified *CUBN* rare variant rs141640975 (GRCh37.p13 Position: 16,992,011, MAF: 0.008) was carried out with respect to the known *CUBN* common SNP rs1801239 (GRCh37.p13 Position: 16,919,052, MAF: 0.10) which is ~73 kbp distant (LD: r^2^:0.0002, D’:1.0) with each other. The effect estimates for the novel variant rs141640975 did not change before *(p*_*rs141640975*_: 8.8 × 10^−7^,β: 0.33) and after conditioning (*P*_*rs141640975_condition*_:8.5 × 10^−7^, β: 0.33) with the known *CUBN* SNP rs1801239 *(P*_*rs1so1239*_*:0.0002*, β: 0.05) as depicted in **Supp. table S1.4.** Conditional Locus zoom plot for *CUBN* index rare variant and known common SNP available in Supp. Material (**Supp. figure S2d**).

### Gene Aggregate Tests

Applying the Hom-O-SKAT Meta (weighted) optimal test on data from six studies (five discovery phase cohorts and the MDCS cohort) comprising a total of 15,867 individuals, we identified three genes surviving Bonferroni correction threshold (*P*_Bon*f*erroni_<2.7 × 10^−6^, n_*genesets*_: 18,026; **Table 3**). These were *HES1* (*p*: 3.7 × 10^−9^) in chromosome 3, *CDC73* (*p*: 6.4 × 10^−9^) in chromosome 1, and *GRM5* (*p*: 1.6 × 10^−6^) in chromosome 11 (**Table 3**). The number of SNPs in each gene ranged between 2 to 7.

**Table 3:** Genes associated with albuminuria through gene-aggregate tests.

| Gene | Chr | P value | N<sub>snp</sub>/set | Full Gene name | Alternate Gene Symbols | Gene_ID (NCBI) Dec, 21, 2017 |
| --- | --- | --- | --- | --- | --- | --- |
| *HES1* | 3q29 | $3.7 \times 10^{-9}$ | 2 | hes family bHLH transcription factor 1 | HHL; HRY; HES-1; bHLHb39 | 3280 |
| *CDC73* | 1q31.2 | $6.4 \times 10^{-9}$ | 2 | cell division cycle 73 | HYX; FIHP; HPTJT; HRPT1; HRPT2; C1orf28 | 79577 |
| *GRM5* | 11q14.2-q14.3 | $1.6 \times 10^{-6}$ | 7 | Glutamate metabotropic receptor 5 | mGlu5; GPRC1E; MGLUR5; PPP1R86 | 2915 |

### SNP-Other trait associations (GWAS, eQTL, Metabolomics)

Pheno Scanner based results on published literature suggests that the *CUBN* rare SNP is associated with creatinine levels in a blood based metabolomics study (*p*: 0.014).

The suggestive *KCNK5* SNP is a known CAD GWAS significant locus. Additional traits associated with rs10947789 C allele include urinary albumin creatinine ratio (UACR, CKD-gen consortia, β: 025, *p*: 5.6 × 10^−4^), myocardial infarction (CARDIoGRAMplusC4D consortia, β: -0.059, *p*: 1.4 × 10^−6^), visual refractive error (β: 0.11, *p*: 0.003), and birthweight (Early Growth Genetics (EGG) Consortium, β: 0.025, *p*<0.007).

eQTL data suggests rs10947789 C allele specific gene expression associations within adrenal gland (β: 0.51, *p*: 2.5 × 10^−5^), subcutaneous adipose tissue (β: 0.34, *p*: 7.2 × 10^−5^), lymphoblastoid cell line (β: 0.03, *p*: 0.0019) and tibial nerve (β: 0.15, *p*: 0.002).

Detailed SNP-trait associations (GWAS, eQTL and serum metabolome) have been documented in **Supp. tables S1.5a-c**.

### SNP Functionality Prediction: SIFT

We used the SIFT (Sorting Intolerant from Tolerant) database (web version) to check whether any top coding variant was associated with functional by nature (http://sift.bii.a-star.edu.sg/). *CUBN* missense rare SNP rs141640975 was described as functionally deleterious with a SIFT median value of 3.44 **(Supp. Table S1.6).**

## Discussion

In the discovery meta Ex-WAS of 13,226 Europeans (5 cohorts) followed by replication in 20,759 Europeans (12 cohorts) and 2,605 Greenlandic Inuit (2 cohorts), we identify one novel *CUBN* variant associated with albuminuria levels in the combined European meta-analyses.

Albeit *CUBN* is a known locus for albuminuria, the identified rare missense variant shows independent effects (with respect to the known *CUBN* common SNP), stronger within the diabetes versus non-diabetes group *(p*_*interacaon*_: 5.4 × 10^−4^). This rare variant explains upto 6.4% variance per rare allele (in a model adjusted for age and gender) of albuminuria levels (natural log transformed). Also the gene-based tests identify three additional genes *(HES1, CDC73*, and *GRM5)* that associate with albuminuria (*p*_Bonferroni_<2.7 × 10^−6^) in a meta-analyses comprising six Scandinavian cohorts. Another SNP passing discovery stage in the *KCNK5* gene replicates (*p* = 0.03) and seems interesting, but does not reach the genome wide threshold in the combined European-Greenlandic (EUR-GL) meta-analyses *(p:* 9.1 × 10^−6^, **Table 2**).

Albeit there have been a few albuminuria GWAS in the past decade^6-8, 16^ they have all examined the common genetic variants (MAF>0.05). On the contrary this study explores the low frequency (0.01> MAF<0.05) and rare (MAF &#x2264; 0.01) variant range particularly from the coding region (exome) of the genome. Some of these rare variants may explain part of the missing heritability.

While common variants in *CUBN* have been previously reported to associate with albuminuria in individuals of European, African, and Hispanic ancestry^6, 17^ the rare missense (A1690V) SNP rs141640975 in *CUBN* which we identify is not in LD with the known lead SNP rs1801239 (r^2^_LD_: 0.0002, D’_LD_- 1.0) for European ancestry. The conditional analyses confirms rs141640975 to be an independent signal with respect to the known common SNP rs1801239 (*p*_conditional:_ 8.5 × 10^−7^, **Supp table S1.4**) with the minor allele (A) associated with increased levels of albuminuria. Further analyses stratified on diabetes status revealed a 3.5 times higher effect of rs141640975 among DM compared to NDM (**Supp. table S1.3**) group in the discovery metaanalyses suggesting potential clinical implications.

Cubilin, encoded by the *CUBN* gene, is expressed in the apical brush border of proximal renal tubule cells (PTCs) and forms a complex with megalin protein to promote albumin re-uptake^6^, ^18^. Since cubilin protein is a co-receptor not only for tubular resorption but also for the intestinal vitamin B_12_-intrinsic factor complex, *CUBN* mutations lead to hereditary form of megaloblastic anemia (or Imerslund-Grasbeck syndrome) characterized by tubular proteinuria and vitamin B12 malabsorption ^21, 23^. Moreover, a recent exome sequencing study revealed a homozygous frameshift mutation in *CUBN* associated with the only cause of proteinuria in affected family members^21^. Despite being a disease gene, recent exome sequencing studies and related reference databases (ExAC^20^) have shown that damaging variants of *CUBN* are rather frequent in “non-diseased” populations and are thus well tolerated by humans^19, 20^. On this basis, it was recently hypothesized that the tubular proteinuria caused by cubilin deficiency could actually be protective against tubular overload, seen, for example, in nephrotic syndrome or even diabetic kidney disease ^19^ As *CUBN* rs141640975 has been shown to be associated with lower serum creatinine (p: 0.014, **Supp. table S1.5c**) in a recent meta-GWAS of circulating metabolites^22^ and causes albuminuria also in the general population group, our study supports the idea that functional variations in *CUBN* might not be damaging and instead protective. The SIFT database testing functionality of the non-synonymous SNPs resulting in amino acid changes (based on sequence homology and the physical properties of amino acids), suggests *CUBN* rs141640975 to be a functional (**Supp. table S1.6**).

The suggestive common SNP rs10947789 in the *KCNK5* locus previously a known locus for CAD^24, 25^ (**Supp. table S1.5a**) encodes the potassium two pore domain channel subfamily K member 5 protein which is mainly expressed in the cortical distal tubules and collecting ducts of the kidney. This protein is highly sensitive to pH^26^, and its functional inactivation may lead to renal acidosis ^27^, dysregulated membrane potential of pulmonary artery myocytes ^28^, and apoptotic volume decrease ^29^, as suggested by animal and *in vitro* studies.

Pheno Scanner data mining resource, revealed rs 10947789 minor allele (C) associated with increased albuminuria (pooled and diabetes subset)^7^, visual refractive error^30^, and birthweight^31^ in Europeans, while lumbar spine bone mineral density^32^ in European and East Asian ancestries (**Supp. table S1.5a)**. This was supportive of what we find in the current study. eQTL based look ups indicate rs10947789 associated strongly with *KCNK5* expression in the adrenal gland *(p:* 2.5 × 10^−5^) and subcutaneous adipose tissue *(p:* 7.2 × 10^−5^), while nominally in the nervous system tissues (p<0.01), suggesting a functional role in the kidneys and adipose tissue (**Supp. table S1.5b**). KCNK5 is also associated with Balkan endemic nephropathy^33^. Overall, the *KCNK5* locus may play an important role in the cardio-renal axis through modification in albuminuria levels.

We performed gene-based tests in order to capture the aggregate effect of gene-specific rare and low frequency variants on albuminuria. Three new genes that were found associated with albuminuria where *HES1, CDC73* and *GRM5.*

*HES1* gene is a transcription factor that is ubiquitously expressed in most organs, also including the kidneys, and has been documented to be involved in Notch signaling pathways that play a role in renal fibrosis^34^, glomerulosclerosis^35^, and other forms of kidney disease^34, 36^. The *CDC73* gene is a tumor suppressor gene whose mutations have been associated to hyperparathyroidism-jaw syndrome and familial hyperparathyroidism^37^. Albuminuria is associated with hyperparathyroidism which is a complication of CKD^38^, and the present findings may thus suggest a plausible link between the two.

*GRM5* encodes glutamate metabotropic receptor 5 which is a G protein coupled receptor involved in second messenger signaling activated by phosphatidylinositol-calcium. Variants in the metabotropic glutamate receptor group I pathway, including *GRM1* and *GRM5*, were strongly enriched in the pathway analysis of a previous GWAS on albuminuria in individuals with type 1 diabetes from the FinnDiane study^8^, without any overlapping individuals with the current gene aggregate meta-analysis. *GRM5* was recently documented to be expressed in podocytes, also associating with podocyte apoptosis in animals^39^, and reported pharmacological effects in humans^40^.

In summary, we identify a novel rare coding *CUBN* variant implicated in elevated albuminuria levels especially in individuals with diabetes. Further we also identify additional novel genes associated with albuminuria through an alternate gene-aggregate approach among Europeans. Our findings provide fresh insights into the genetic architecture of albuminuria and highlight new targets, genes and pathways for the prevention and treatment of diabetic kidney disease.

## Author Contributions

T.S.A. and T.H. designed the study. T.S.A. was responsible for the overall data collection, and data management. T.H., N.G., A.L., T.S., B.T., M.J., I.B., C.K.C. were responsible for subject recruitment, cohort management and genotyping of the discovery set cohorts along with the DanFUNd, and Greenlanders study. T.S.A. drafted and revised the paper. T.S.A. made the figures and analyzed all data in the discovery set and for the DanFUNd cohort in replication set and also performed meta-analyses at all levels. T.S.A., and J.W., analyzed the gene aggregate tests. C.A.S., and P.A. performed replication and gene aggregate testing for MDCS cohort. R.C. performed replication for Genesis/Gendiab cohort. N.S., N.V.Z, and E.A. performed replication for SUMMIT Consortia cohorts. N.G. performed replication for Greenlanders cohort. T.S.A., T.O.K., N.G., N.S., P-H.G, M.S., L.G., M.M., O.M., M.O-M., P.R. and T.H. were responsible for the data interpretation and useful comments/suggestions. All authors reviewed and approved the final version of the manuscript.

## Acknowledgements and Funding

Study specific acknowledgements and funding details available under the **Supplementary Material Supp. table S1.7.** SUMMIT consortia members listed in Supp. Table S1.8.

T.S.A. was supported by Lundbeck Foundation travel grant Reference #2013-14471. I thank Puneet Gandhi for her helpful feedback on the study from a layman’s point of view.

## Disclosure Statements

P-H.G. has received investigator-initiated research grants from Eli Lilly and Roche, is an advisory board member for AbbVie, AstraZeneca, Boehringer Ingelheim, Cebix, Eli Lilly, Janssen, Medscape, Merck Sharp & Dohme, Novartis, Novo Nordisk and Sanofi; and has received lecture fees from AstraZeneca, Boehringer Ingelheim, Eli Lilly, Elo Water, Genzyme, Merck Sharp & Dohme, Medscape, Novo Nordisk and Sanofi.

## References

1. National Institutes of Health, NIDDK, Bethesda, MD.: United States Renal Data System. 2017 USRDS annual data report: Epidemiology of kidney disease in the United States., 2017.

2. Carrero JJ, Grams ME, Sang Y, Arnlov J, Gasparini A, Matsushita K, Qureshi AR, Evans M, Barany P, Lindholm B, Ballew SH, Levey AS, Gansevoort RT, Eiinder CG, Coresh J: Albuminuria changes are associated with subsequent risk of end-stage renal disease and mortality. Kidney international, 91: 244–251, 2017.

3. Skaaby T, Husemoen LL, Ahluwalia TS, Rossing P, Jorgensen T, Thuesen BH, Pisinger C, Rasmussen K, Linneberg A: Cause-specific mortality according to urine albumin creatinine ratio in the general population. PloS one, 9: e93212, 2014.

4. Ishigami J, Grams ME, Naik RP, Caughey MC, Loehr LR, Uchida S, Coresh J, Matsushita K: Hemoglobin, Albuminuria, and Kidney Function in Cardiovascular Risk: The ARIC (Atherosclerosis Risk in Communities) Study. J Am Heart Assoc, 7, 2018.

5. Nichols GA, Deruaz-Luyet, A, Hauske SJ, Brodovicz, KG: The association between estimated glomerular filtration rate, albuminuria, and risk of cardiovascular hospitalizations and all-cause mortality among patients with type 2 diabetes. J Diabetes Complications, 32: 291–297, 2018.

6. Boger CA, Chen MH, Tin A, Olden M, Kottgen A, de Boer, IH, Fuchsberger C, O’Seaghdha, CM, Pattaro, Teumer, A, Liu, CT, Glazer, NL, Li, M, O’Connell, JR, Tanaka, T, Peralta, CA, Kutalik, Z, Luan, J, Zhao, JH, Hwang, SJ, Akylbekova, E, Kramer, H, van der Harst, P, Smith, AV, Lohman, K, de Andrade, M, Hayward, C, Kollerits, B, Tonjes, A, Aspelund, T, Ingelsson, E, Eiriksdottir, G, Launer LJ, Harris TB, Shuldiner AR, Mitchell BD, Arking DE, Franceschini N, Boerwinkle E, Egan J, Hernandez D, Reilly M, Townsend RR, Lumley T, Siscovick DS, Psaty, BM, Kestenbaum, B, Haritunians, T, Bergmann, S, Vollenweider, P, Waeber, G, Mooser, V, Waterworth, D, Johnson, AD, Florez, JC, Meigs, JB, Lu, X, Turner, ST, Atkinson, EJ, Leak, TS, Aasarod, K, Skorpen, F, Syvanen, AC, Illig, T, Baumert, J, Koenig, W, Kramer, BK, Devuyst, O, Mychaleckyj, JC, Minelli, C, Bakker, SJ, Kedenko, L, Paulweber, B, Coassin, S, Endlich, K, Kroemer, HK, Biffar, R, Stracke, S, Volzke, H, Stumvoll, M, Magi, R, Campbell, H, Vitart, V, Hastie, ND, Gudnason, V, Kardia, SL, Liu, Y, Polasek, O, Curhan, G, Kronenberg, F, Prokopenko, I, Rudan, I, Arnlov, J, Hallan, S, Navis, G, Consortium, CK, Parsa, A, Ferrucci, L, Coresh, J, Shlipak, MG, Bull, SB, Paterson, NJ, Wichmann, HE, Wareham, NJ, Loos, RJ, Rotter, JI, Pramstaller, PP, Cupples, LA, Beckmann, JS, Yang, Q, Heid, IM, Rettig, R, Dreisbach, AW, Bochud, M, Fox, CS, Kao, WH: CUBN is a gene locus for albuminuria. Journal of the American Society of Nephrology: JASN, 22: 555–570, 2011.

7. Teumer A, Tin A, Sorice R, Gorski M, Yeo NC, Chu AY, Li M, Li Y, Mijatovic V, Ko YA, Taliun D, Luciani A, Chen MH, Yang Q, Foster MC, Olden M, Hiraki LT, Tayo BO, Fuchsberger C, Dieffenbach AK, Shuldiner AR, Smith AV, Zappa AM, Lupo A, Kollerits B, Ponte B, Stengel B, Kramer BK, Paulweber B, Mitchell BD, Hayward C, Helmer C, Meisinger C, Gieger C, Shaffer CM, Muller C, Langenberg C, Ackermann D, Siscovick D, Dcct/Edic, Boerwinkle, E, Kronenberg, F, Ehret, GB, Homuth, G, Waeber, G, Navis, G, Gambaro, G, Malerba, G, Eiriksdottir, G, Li, G, Wichmann, HE, Grallert, H, Wallaschofski, H, Volzke, H, Brenner, H, Kramer, H, Mateo Leach, I, Rudan, I, Hillege, HL, Beckmann, JS, Lambert, JC, Luan, J, Zhao, JH, Chalmers, J, Coresh, J, Denny, JC, Butterbach, K, Launer, LJ, Ferrucci, L, Kedenko, L, Haun, M, Metzger, M, Woodward, M, Hoffman, MJ, Nauck, M, Waldenberger, M, Pruijm, M, Bochud, M, Rheinberger, M, Verweij, N, Wareham, NJ, Endlich, N, Soranzo, N, Polasek, O, van der Harst, P, Pramstaller, PP, Vollenweider, P, Wild, PS, Gansevoort, RT, Rettig, R, Biffar, R, Carroll, RJ, Katz, R, Loos, RJ, Hwang, SJ, Coassin, S, Bergmann, S, Rosas, SE, Stracke, S, Harris, TB, Corre, T, Zeller, T, Illig, T, Aspelund, T, Tanaka, T, Lendeckel, U, Volker, U, Gudnason, V, Chouraki, V, Koenig, W, Kutalik, Z, O’Connell, JR, Parsa, A, Heid, IM, Paterson, AD, de Boer, IH, Devuyst, O, Lazar, J, Endlich, K, Susztak, K, Tremblay, J, Hamet, P, Jacob, HJ, Boger, CA, Fox, CS, Pattaro, C, Kottgen, A: Genome-wide Association Studies Identify Genetic Loci Associated With Albuminuria in Diabetes. Diabetes, 65: 803–817, 2016.

8. Sandholm N, Forsblom C, Makinen VP, McKnight AJ, Osterholm AM, He B, Harjutsalo V, Lithovius R, Gordin D, Parkkonen M, Saraheimo M, Thorn LM, Tolonen N, Waden J, Tuomilehto J, Lajer M, Ahlqvist E, Mollsten A, Marcovecchio ML, Cooper J, Dünger, D, Paterson, AD, Zerbini, G, Groop, L, Consortium, S, Tarnow, L, Maxwell, AP, Tryggvason, K, Groop, PH, FinnDiane Study, G: Genome-wide association study of urinary albumin excretion rate in patients with type 1 diabetes. Diabetologia, 57: 1143–1153, 2014.

9. Li M, Li Y, Weeks, O, Mijatovic, V, Teumer, A, Huffman, JE, Tromp, G, Fuchsberger, C, Gorski, M, Lyytikainen, LP, Nutile, T, Sedaghat, S, Sorice, R, Tin, A, Yang, Q, Ahluwalia, TS, Arking, DE, Bihlmeyer, NA, Boger, CA, Carroll, RJ, Chasman, DI, Cornelis, MC, Dehghan, A, Faul, JD, Feitosa, MF, Gambaro, G, Gasparini, P, Giulianini, F, Heid, I, Huang, J, Imboden, M, Jackson, AU, Jeff, J, Jhun, MA, Katz, R, Kifley, A, Kilpelainen, TO, Kumar, A, Laakso, M, Li-Gao, R, Lohman, K, Lu, Y, Magi, R, Malerba, G, Mihailov, E, Mohlke, KL, Mook-Kanamori, DO, Robino, A, Ruderfer, D, Salvi, E, Schick, UM, Schulz, CA, Smith, AV, Smith, JA, Traglia, M, Yerges-Armstrong, LM, Zhao, W, Goodarzi, MO, Kraja, AT, Liu, C, Wessel, J, Group, CG-TDW, Group, CBPW, Boerwinkle, E, Borecki, IB, Bork-Jensen, J, Bottinger, EP, Braga, D, Brandslund, I, Brody, JA, Campbell, A, Carey, DJ, Christensen, C, Coresh, J, Crook, E, Curhan, GC, Cusi, D, de Boer, IH, de Vries, AP, Denny, JC, Devuyst, O, Dreisbach, AW, Endlich, K, Esko, T, Franco, OH, Fulop, T, Gerhard, GS, Glumer, C, Gottesman, O, Grarup, N, Gudnason, V, Hansen, T, Harris, TB, Hayward, C, Hocking, L, Hofman, A, Hu, FB, Husemoen, LL, Jackson, RD, Jorgensen, T, Jorgensen, ME, Kahonen, M, Kardia, SL, König, W, Kooperberg, C, Kriebel, J, Launer, LJ, Lauritzen, T, Lehtimaki, T, Levy, D, Linksted, P, Linneberg, A, Liu, Y, Loos, RJ, Lupo, A, Meisinger, C, Melander, O, Metspalu, A, Mitchell, P, Nauck, M, Nürnberg, P, Orho-Melander, M, Parsa, A, Pedersen, O, Peters, A, Peters, U, Polasek, O, Porteous, D, Probst-Hensch, NM, Psaty, BM, Qi, L, Raitakari, OT, Reiner, AP, Rettig, R, Ridker, PM, Rivadeneira, F, Rossouw, JE, Schmidt, F, Siscovick, D, Soranzo, N, Strauch, K, Toniolo, D, Turner, ST, Uitterlinden, AG, Ulivi, S, Velayutham, D, Volker, U, Volzke, H, Waldenberger, M, Wang, JJ, Weir, DR, Witte, D, Kuivaniemi, H, Fox, CS, Franceschini, N, Goessling, W, Kottgen, A, Chu, AY: S0S2 and ACPI Loci Identified through Large-Scale Exome Chip Analysis Regulate Kidney Development and Function. Journal of the American Society of Nephrology: JASN, 28: 981–994, 2017.

10. Ahluwalia TS, Allin KH, Sandholt CH, Sparso TH, Jorgensen ME, Rowe M, Christensen C, Brandslund Lauritzen, T Linneberg, A Husemoen, LL Jorgensen, T Hansen, T Grarup, N Pedersen, O: Discovery of coding genetic variants influencing diabetes-related serum biomarkers and their impact on risk of type 2 diabetes. J Clin Endocrinol Metab, 100: E664–671, 2015.

11. Ahluwalia TS, Troelsen JT, Balslev-Harder, M, Bork-Jensen, J, Thuesen, BH, Cerqueira, C, Linneberg, A, Grarup, N, Pedersen, O, Hansen, T, Dalgaard, LT: Carriers of a VEGFA enhancer polymorphism selectively binding CHOP/DDIT3 are predisposed to increased circulating levels of thyroid-stimulating hormone. J Med Genet, 54: 166–175, 2017.

12. van Zuydam, NR Ahlqvist, E Sandholm, N Deshmukh, H Rayner, NW Abdalla, M Ladenvall, C Ziemek, D Fauman, E Robertson, NR McKeigue, PM Valo, E Forsblom, C Harjutsalo, V centres, FS Perna, A Rurali, E Marcovecchio, ML Igo, RP, Jr., Salem, RM, Perico, N, Lajer, M, Karajamaki, A, Imamura, M, Kubo, M, Takahashi, A, Sim, X, Liu, J, van Dam, RM, Jiang, G, Tam, CHT, Luk, AOY, Lee, HM, Lim, CKP, Szeto, CC, So, WY, Chan, JCN, Hong Kong Diabetes Registry, TRSPG, Ang, SF, Dorajoo, R, Wang, L, Hua Clara, TS, McKnight, AJ, Duffy, S, Warren, UKGSG, Pezzolesi, MG, Consortium, G, Marre, M, Gyorgy, B, Hadjadj, S, Hiraki, LT, group, DE, Ahluwalia, TS, Almgren, P, Schulz, CA, Orho-Melander, M, Linneberg, A, Christensen, C, Witte, DR, Grarup, N, Brandslund, I, Melander, O, Paterson, AD, Tregouet, D, Maxwell, AP, Lim, SC, Ma, RCW, Tai, ES, Maeda, S, Lyssenko, V, Tuomi, T, Krolewski, AS, Rich, SS, Hirschhorn, JN, Florez, JC, Dünger, D, Pedersen, O, Hansen, T, Rossing, P, Remuzzi, G, Consortium, S, Brosnan, MJ, Palmer, CNA, Groop, PH, Colhoun, HM, Groop, LC, McCarthy, MI: A Genome-Wide Association Study of Diabetic Kidney Disease in subjects With Type 2 Diabetes. Diabetes, 2018.

13. Albrechtsen A, Grarup N, Li Y, Sparso T, Tian G, Cao H, Jiang T, Kim SY, Korneliussen T, Li Q, Nie, C, Wu R, Skotte L, Morris AP, Ladenvall C, Cauchi S, Stancakova A, Andersen G, Astrup A, Banasik K, Bennett AJ, Bolund L, Charpentier G, Chen Y, Dekker JM, Doney AS, Dorkhan M, Forsen T, Frayling TM, Groves CJ, Gui Y, Hallmans G, Hattersley AT, He K, Hitman GA, Holmkvist J, Huang S, Jiang H, Jin X, Justesen JM, Kristiansen K, Kuusisto J, Lajer M, Lantieri, O, Li, W, Liang, H, Liao, Q, Liu, X, Ma, T, Ma, X, Manijak, MP, Marre, M, Mokrosinski, J, Morris, AD, Mu, B, Nielsen, AA, Nijpels, G, Nilsson, P, Palmer, CN, Rayner, NW, Renstrom, F, Ribel-Madsen, R, Robertson, N, Rolandsson, O, Rossing, P, Schwartz, TW, Group, DESIRS, Slagboom, PE, Sterner, M, Consortium, D, Tang, M, Tarnow, L, Tuomi, T, van’t Riet, E, van Leeuwen, N, Varga, TV, Vestmar, MA, Walker, M, Wang, B, Wang, Y, Wu, H, Xi, F, Yengo, L, Yu, C, Zhang, X, Zhang, J, Zhang, Q, Zhang, W, Zheng, H, Zhou, Y, Altshuler, D, t Hart, LM, Franks, PW, Balkau, B, Froguel, P, McCarthy, MI, Laakso, M, Groop, L, Christensen, C, Brandslund, I, Lauritzen, T, Witte, DR, Linneberg, A, Jorgensen, T, Hansen, T, Wang, J, Nielsen, R, Pedersen, O: Exome sequencing-driven discovery of coding polymorphisms associated with common metabolic phenotypes. Diabetologia, 56: 298–310, 2013.

14. Lee S, Teslovich TM, Boehnke M, Lin X: General framework for meta-analysis of rare variants in sequencing association studies. Am J Hum Genet, 93: 42–53, 2013.

15. Staley JR, Blackshaw J, Kamat MA, Ellis S, Surendran P, Sun BB, Paul DS, Freitag D, Burgess S, Danesh J, Young R, Butterworth AS: PhenoScanner: a database of human genotype-phenotype associations. Bioinformatics, 32: 3207–3209, 2016.

16. Pattaro C: Genome-wide association studies of albuminuria: towards genetic stratification in diabetes? J Nephrol, 2017.

17. Kramer HJ, Stilp AM, Laurie CC, Reiner AP, Lash J, Daviglus ML, Rosas SE, Ricardo AC, Tayo BO, Flessner MF, Kerr KF, Peralta C, Durazo-Arvizu, R, Conomos, M, Thornton, T, Rotter, J, Taylor, KD, Cai, J, Eckfeldt, J, Chen, H, Papanicolau, G, Franceschini, N: African Ancestry-Specific Alleles and Kidney Disease Risk in Hispanics/Latinos. Journal of the American Society of Nephrology: JASN, 28: 915–922, 2017.

18. Amsellem S, Gburek J, Hamard G, Nielsen R, Willnow TE, Devuyst, O, Nexo, E, Verroust, PJ, Christensen, EI, Kozyraki, R: Cubilin is essential for albumin reabsorption in the renal proximal tubule. Journal of the American Society of Nephrology: JASN, 21:1859–1867, 2010.

19. Simons M: The Benefits of Tubular Proteinuria: An Evolutionary Perspective. Journal of the American Society of Nephrology: JASN, 29: 710–712, 2018.

20. Lek M, Karczewski KJ, Minikel EV, Samocha KE, Banks E, Fennell T, O’Donnell-Luria, AH, Ware, JS, Hill AJ, Cummings BB, Tukiainen T, Birnbaum DP, Kosmicki JA, Duncan LE, Estrada K, Zhao F, Zou J, Pierce-Hoffman, E, Berghout, J, Cooper, DN, Deflaux, N, DePristo, M, Do, R, Flannick, J, Fromer, M, Gauthier, L, Goldstein, J, Gupta, N, Howrigan, D, Kiezun, A, Kurki, MI, Moonshine, AL, Natarajan, P, Orozco, L, Peloso, GM, Poplin, R, Rivas, MA, Ruano-Rubio, V, Rose, SA, Ruderfer, DM, Shakir, K, Stenson, PD, Stevens, C, Thomas, BP, Tiao, G, Tusie-Luna, MT, Weisburd, B, Won, HH, Yu, Altshuler, DM, Ardissino, D, Boehnke, M, Danesh, J, Donnelly, S, Elosua, R, Florez, JC, Gabriel, SB, Getz, G, Glatt, SJ, Hultman, CM, Kathiresan, S, Laakso, M, McCarroll, S, McCarthy, MI, McGovern, D, McPherson, R, Neale, BM, Palotie, A, Purcell, SM, Saleheen, D, Scharf, JM, Sklar, P, Sullivan, PF, Tuomilehto, J, Tsuang, MT, Watkins, HC, Wilson, JG, Daly, MJ, MacArthur, DG, Exome Aggregation, C: Analysis of protein-coding genetic variation in 60,706 humans. Nature, 536: 285–291, 2016.

21. Ovunc B, Otto EA, Vega-Warner, V, Saisawat, P, Ashraf, S, Ramaswami, G, Fathy, HM, Schoeb, D, Chernin, G, Lyons, RH, Yilmaz, E, Hildebrandt, F: Exome sequencing reveals cubilin mutation as a single-gene cause of proteinuria. Journal of the American Society of Nephrology: JASN, 22: 1815–1820, 2011.

22. Kettunen J, Demirkan A, Wurtz P, Draisma HH, Haller T, Rawal R, Vaarhorst A, Kangas AJ, Lyytikainen LP, Pirinen M, Pool R, Sarin AP, Soininen P, Tukiainen T, Wang Q, Tiainen M, Tynkkynen T, Amin N, Zeller T, Beekman M, Deelen J, van Dijk, KW, Esko, T, Hottenga, JJ, van Leeuwen, EM, Lehtimaki, T, Mihailov, E, Rose, RJ, de Craen, AJ, Gieger, C, Kahonen, M, Perola, M, Blankenberg, S, Savolainen, MJ, Verhoeven, A, Viikari, J, Willemsen, G, Boomsma, DI, van Duijn, CM, Eriksson, J, Jula, A, Jarvelin, MR, Kaprio, J, Metspalu, A, Raitakari, O, Salomaa, V, Slagboom, PE, Waldenberger, M, Ripatti, S, Ala-Korpela, M: Genome-wide study for circulating metabolites identifies 62 loci and reveals novel systemic effects of LPA. Nature communications, 7: 11122, 2016.

23. Grasbeck R: Imerslund-Grasbeck syndrome (selective vitamin B(12) malabsorption with proteinuria). Orphanet J Rare Dis, 1:17, 2006.

24. Consortium CAD, Deloukas P, Kanoni S, Willenborg C, Farrall M, Assimes TL, Thompson JR, Ingelsson E, Saleheen D, Erdmann J, Goldstein BA, Stirrups K, König, IR, Cazier, JB, Johansson, A, Hall, AS, Lee, JY, Willer, CJ, Chambers, JC, Esko, T, Folkersen, L, Goel, A, Grundberg, E, Havulinna, AS, Ho, WK, Hopewell, JC, Eriksson, N, Kleber, ME, Kristiansson, K, Lundmark, P, Lyytikainen, LP, Rafelt, S, Shungin, D, Strawbridge, RJ, Thorleifsson, G, Tikkanen, E, Van Zuydam, N, Voight, BF, Waite, LL, Zhang, W, Ziegler, A, Absher, D, Altshuler, D, Balmforth, AJ, Barroso, I, Braund, PS, Burgdorf, C, Claudi-Boehm, S, Cox, D, Dimitriou, M, Do, R, Consortium, D, Consortium, C, Doney, AS, El Mokhtari, N, Eriksson, P, Fischer, K, Fontanillas, P, Franco-Cereceda, A, Gigante, B, Groop, L, Gustafsson, S, Hager, J, Hallmans, G, Han, BG, Hunt, SE, Kang, HM, Illig, T, Kessler, T, Knowles, JW, Kolovou, G, Kuusisto, J, Langenberg, C, Langford, C, Leander, K, Lokki, ML, Lundmark, A, McCarthy, MI, Meisinger, C, Melander, O, Mihailov, E, Maouche, S, Morris, AD, Muller-Nurasyid, M, Mu, TC, Nikus, K, Peden, JF, Rayner, NW, Rasheed, A, Rosinger, S, Rubin, D, Rumpf, MP, Schafer, A, Sivananthan, M, Song, C, Stewart, AF, Tan, ST, Thorgeirsson, G, van der Schoot, CE, Wagner, PJ, Wellcome Trust Case Control, C, Wells, GA, Wild, PS, Yang, TP, Amouyel, P, Arveiler, D, Basart, H, Boehnke, M, Boerwinkle, E, Brambilla, P, Cambien, F, Cupples, AL, de Faire, U, Dehghan, A, Diemert, P, Epstein, SE, Evans, A, Ferrario, MM, Ferrieres, J, Gauguier, D, Go, AS, Goodall, AH, Gudnason, V, Hazen, SL, Holm, H, Iribarren, C, Jang, Y, Kahonen, M, Kee, F, Kim, HS, Klopp, N, Koenig, W, Kratzer, W, Kuulasmaa, K, Laakso, M, Laaksonen, R, Lee, JY, Lind, L, Ouwehand, WH, Parish, S, Park, JE, Pedersen, NL, Peters, A, Quertermous, T, Rader, DJ, Salomaa, V, Schadt, E, Shah, SH, Sinisalo, J, Stark, K, Stefansson, K, Tregouet, DA, Virtamo, J, Wallentin, L, Wareham, N, Zimmermann, ME, Nieminen, MS, Hengstenberg, C, Sandhu, MS, Pastinen, T, Syvanen, AC, Hovingh, GK, Dedoussis, G, Franks, PW, Lehtimaki, T, Metspalu, A, Zalloua, PA, Siegbahn, A, Schreiber, S, Ripatti, S, Blankenberg, SS, Perola, M, Clarke, R, Boehm, BO, O’Donnell, C, Reilly, MP, Marz, W, Collins, R, Kathiresan, S, Hamsten, A, Kooner, JS, Thorsteinsdottir, U, Danesh, J, Palmer, CN, Roberts, R, Watkins, H, Schunkert, H, Samani, NJ: Large-scale association analysis identifies new risk loci for coronary artery disease. Nature genetics, 45: 25–33, 2013.

25. Howson JMM, Zhao W, Barnes DR, Ho WK, Young R, Paul DS, Waite LL, Freitag DF, Fauman EB, Salfati EL, Sun BB, Eicher JD, Johnson AD, Sheu WHH, Nielsen SF, Lin WY, Surendran P, Malarstig A, Wilk JB, Tybjaerg-Hansen, A, Rasmussen, KL, Kamstrup, PR, Deloukas, P, Erdmann, J, Kathiresan, S, Samani, NJ, Schunkert, H, Watkins, H, CardioGramplusC4D, Do, R, Rader, DJ, Johnson, JA, Hazen, SL, Quyyumi, AA, Spertus, JA, Pepine, CJ, Franceschini, N, Justice, A, Reiner, AP, Buyske, S, Hindorff, LA, Carty, CL, North, KE, Kooperberg, C, Boerwinkle, E, Young, K, Graff, M, Peters, U, Absher, D, Hsiung, CA, Lee, WJ, Taylor, KD, Chen, YH, Lee, IT, Guo, X, Chung, RH, Hung, YJ, Rotter, Jl, Juang, JJ, Quertermous, T, Wang, TD, Rasheed, A, Frossard, P, Alam, DS, Majumder, AAS, Di Angelantonio, E, Chowdhury, R, Epic, CVD, Chen, YI, Nordestgaard, BG, Assimes, TL, Danesh, J, Butterworth, AS, Saleheen, D: Fifteen new risk loci for coronary artery disease highlight arterial-wall-specific mechanisms. Nature genetics, 49: 1113–1119, 2017.

26. Morton MJ, Abohamed A, Sivaprasadarao A, Hunter M: pH sensing in the two-pore domain K+ channel, TASK2. Proc Natl Acad Sci U S A, 102: 16102–16106, 2005.

27. Warth R, Barriere H, Meneton P, Bloch M, Thomas J, Taue M, Heitzmann D, Romeo E, Verrey F, Mengual R, Guy N, Bendahhou S, Lesage F, Poujeol P, Barhanin J: Proximal renal tubular acidosis in TASK2 K+ channel-deficient mice reveals a mechanism for stabilizing bicarbonate transport. Proc Natl Acad Sei USA, 101: 8215–8220, 2004.

28. Gonczi M, Szentandrassy N, Johnson IT, Heagerty AM, Weston AH: Investigation of the role of TASK-2 channels in rat pulmonary arteries; pharmacological and functional studies following RNA interference procedures. Br J Pharmacol, 147: 496–505, 2006.

29. L’Hoste, S, Poet, M, Duranton, C, Belfodil, R, e Barriere, H, Rubera, I, Tauc, M, Poujeol, C, Barhanin, J, Poujeol, P: Role of TASK2 in the control of apoptotic volume decrease in proximal kidney cells. J Biol Chem, 282: 36692–36703, 2007.

30. Stambolian D, Wojciechowski R, Oexle K, Pirastu M, Li X, Raffel LJ, Cotch MF, Chew EY, Klein B, Klein R, Wong TY, Simpson CL, Klaver CC, van Duijn, CM, Verhoeven, VJ, Baird, PN, Vitart, V, Paterson, AD, Mitchell, P, Saw, SM, Fossarello, M, Kazmierkiewicz, K, Murgia, F, Portas, L, Schache, M, Richardson, A, Xie, J, Wang, JJ, Rochtchina, E, Group, DER, Viswanathan, AC, Hayward, C, Wright, AF, Polasek, O, Campbell, H, Rudan, I, Oostra, BA, Uitterlinden, AG, Hofman, A, Rivadeneira, F, Amin, N, Karssen, LC, Vingerling, JR, Hosseini, SM, Doring, A, Bettecken, T, Vatavuk, Z, Gieger, C, Wichmann, HE, Wilson, JF, Fleck, B, Foster, PJ, Topouzis, F, McGuffin, P, Sim, X, Inouye, M, Holliday, EG, Attia, J, Scott, RJ, Rotter, JI, Meitinger, T, Bailey-Wilson, JE: Meta-analysis of genome-wide association studies in five cohorts reveals common variants in RBFOX1, a regulator of tissue-specific splicing, associated with refractive error. Human molecular genetics, 22: 2754–2764, 2013.

31. Horikoshi M, Yaghootkar H, Mook-Kanamori, DO, Sovio, U, Taal, HR, Hennig, BJ, Bradfield, JP, St Pourcain, B, Evans, DM, Charoen, P, Kaakinen, M, Cousminer, DL, Lehtimaki, T, Kreiner-Moller, E, Warrington, NM, Bustamante, M, Feenstra, B, Berry, DJ, Thiering, E, Pfab, T, Barton, SJ, Shields, BM, Kerkhof, M, van Leeuwen, EM, Fulford, AJ, Kutalik, Z, Zhao, JH, den Hoed, M, Mahajan, A, Lindi, V, Goh, LK, Hottenga, JJ, Wu, Y, Raitakari, OT, Harder, MN, Meirhaeghe, A, Ntalla, I, Salem, RM, Jameson, KA, Zhou, K, Monies, DM, Lagou, V, Kirin, M, Heikkinen, J, Adair, LS, Alkuraya, FS, Al-Odaib, A, Amouyel, P, Andersson, EA, Bennett, AJ, Blakemore, AI, Buxton, JL, Dallongeville, J, Das, S, de Geus, EJ, Estivill, X, Flexeder, C, Froguel, P, Geller, F, Godfrey, KM, Gottrand, F, Groves, CJ, Hansen, T, Hirschhorn, JN, Hofman, A, Hollegaard, MV, Hougaard, DM, Hypponen, E, Inskip, HM, Isaacs, A, Jorgensen, T, Kanaka-Gantenbein, C, Kemp, JP, Kiess, W, Kilpelainen, TO, Klopp, N, Knight, BA, Kuzawa, CW, McMahon, G, Newnham, JP, Niinikoski, H, Oostra, BA, Pedersen, L, Postma, DS, Ring, SM, Rivadeneira, F, Robertson, NR, Sebert, S, Simell, O, Slowinski, T, Tiesler, CM, Tonjes, A, Vaag, A, Viikari, JS, Vink, JM, Vissing, NH, Wareham, NJ, Willemsen, G, Witte, DR, Zhang, H, Zhao, J, Meta-Analyses of, G, Insulin-related traits, C, Wilson, JF, Stumvoll, M, Prentice, AM, Meyer, BF, Pearson, ER, Boreham, CA, Cooper, C, Gillman, MW, Dedoussis, GV, Moreno, LA, Pedersen, O, Saarinen, M, Mohlke, KL, Boomsma, DI, Saw, SM, Lakka, TA, Korner, A, Loos, RJ, Ong, KK, Vollenweider, P, van Duijn, CM, Koppelman, GH, Hattersley, AT, Holloway, JW, Hocher, B, Heinrich, J, Power, C, Melbye, M, Guxens, M, Pennell, CE, Bonnelykke, K, Bisgaard, H, Eriksson, JG, Widen, E, Hakonarson, H, Uitterlinden, AG, Pouta, A, Lawlor, DA, Smith, GD, Frayling, TM, McCarthy, MI, Grant, SF, Jaddoe, VW, Jarvelin, MR, Timpson, NJ, Prokopenko, I, Freathy, RM, Early Growth Genetics, C: New loci associated with birth weight identify genetic links between intrauterine growth and adult height and metabolism. Nature genetics, 45: 76–82, 2013.

32. Estrada K, Styrkarsdottir U, Evangelou E, Hsu YH, Duncan EL, Ntzani EE, Oei L, Albagha OM, Amin N, Kemp JP, Koller DL, Li G, Liu CT, Minster RL, Moayyeri A, Vandenput L, Willner D, Xiao SM, Yerges-Armstrong, LM, Zheng, HF, Alonso, N, Eriksson, J, Kammerer, CM, Kaptoge, SK, Leo, PJ, Thorleifsson, G, Wilson, SG, Wilson, JF, Aalto, V, Alen, M, Aragaki, AK, Aspelund, T, Center, JR, Dailiana, Z, Duggan, DJ, Garcia, M, Garcia-Giralt, N, Giroux, S, Hallmans, G, Hocking, U, Husted, LB, Jameson, KA, Khusainova, R, Kim, GS, Kooperberg, C, Koromila, T, Kruk, M, Laaksonen, M, Lacroix, AZ, Lee, SH, Leung, PC, Lewis, JR, Masi, L, Mencej-Bedrac, S, Nguyen, TV, Nogues, X, Patel, MS, Prezelj, J, Rose, LM, Scollen, S, Siggeirsdottir, K, Smith, AV, Svensson, O, Trompet, S, Trümmer, O, van Schoor, NM, Woo, J, Zhu, K, Balcells, S, Brandi, ML, Buckley, BM, Cheng, S, Christiansen, C, Cooper, C, Dedoussis, G, Ford, I, Frost, M, Goltzman, D, Gonzalez-Macias, J, Kahonen, M, Karlsson, M, Khusnutdinova, E, Koh, JM, Kollia, P, Langdahl, BL, Leslie, WD, Lips, P, Ljunggren, O, Lorenc, RS, Marc, J, Mellstrom, D, Obermayer-Pietsch, B, Olmos, JM, Pettersson-Kymmer, U, Reid, DM, Riancho, JA, Ridker, PM, Rousseau, F, Slagboom, PE, Tang, NL, Urreizti, R, Van Hul, W, Viikari, J, Zarrabeitia, MT, Aulchenko, YS, Castano-Betancourt, M, Grundberg, E, Herrera, L, Ingvarsson, T, Johannsdottir, H, Kwan, T, Li, R, Luben, R, Medina-Gomez, C, Palsson, ST, Reppe, S, Rotter, JI, Sigurdsson, G, van Meurs, JB, Verlaan, D, Williams, FM, Wood, AR, Zhou, Y, Gautvik, KM, Pastinen, T, Raychaudhuri, S, Cauley, JA, Chasman, DI, Clark, GR, Cummings, SR, Danoy, P, Dennison, EM, Eastell, R, Eisman, JA, Gudnason, V, Hofman, A, Jackson, RD, Jones, G, Jukema, JW, Khaw, KT, Lehtimaki, T, Liu, Y, Lorentzon, M, McCloskey, E, Mitchell, BD, Nandakumar, K, Nicholson, GC, Oostra, BA, Peacock, M, Pols, HA, Prince, RL, Raitakari, O, Reid, IR, Robbins, J, Sambrook, PN, Sham, PC, Shuldiner, AR, Tylavsky, FA, van Duijn, CM, Wareham, NJ, Cupples, LA, Econs, MJ, Evans, DM, Harris, TB, Kung, AW, Psaty, BM, Reeve, J, Spector, TD, Streeten, EA, Zillikens, MC, Thorsteinsdottir, U, Ohlsson, C, Karasik, D, Richards, JB, Brown, MA, Stefansson, K, Uitterlinden, AG, Ralston, SH, Ioannidis, JP, Kiel, DP, Rivadeneira, F: Genome-wide meta-analysis identifies 56 bone mineral density loci and reveals 14 loci associated with risk of fracture. Nature genetics, 44: 491–501, 2012.

33. Toncheva D, Mihailova-Hristova, M, Vazharova, R, Staneva, R, Karachanak, S, Dimitrov, P, Simeonov, V, Ivanov, S, Balabanski, L, Serbezov, D, Malinov, M, Stefanovic, V, Cukuranovic, R, Polenakovic, M, Jankovic-Velickovic, L, Djordjevic, V, Jevtovic-Stoimenov, T, Plaseska-Karanfilska, D, Galabov, A, Djonov, V, Dimova, I: NGS nominated CELA1, HSPG2, and KCNK5 as candidate genes for predisposition to Balkan endemic nephropathy. Biomed Res Int, 2014: 920723, 2014.

34. Huang R, Zhou Q, Veeraragoo P, Yu H, Xiao Z: Notch2/Hes-1 pathway plays an important role in renal ischemia and reperfusion injury-associated inflammation and apoptosis and the gamma-secretase inhibitor DAPT has a nephroprotective effect. Ren Fail, 33: 207–216, 2011.

35. Ueno T, Kobayashi N, Nakayama M, Takashima Y, Ohse T, Pastan I, Pippin JW, Shankland SJ, Uesugi N, Matsusaka T, Nagata M: Aberrant Notch1-dependent effects on glomerular parietal epithelial cells promotes collapsing focal segmental glomerulosclerosis with progressive podocyte loss. Kidney international, 83: 1065–1075, 2013.

36. Kobayashi T, Terada Y, Kuwana H, Tanaka H, Okado T, Kuwahara M, Tohda S, Sakano S, Sasaki S: Expression and function of the Delta-1/Notch-2/Hes-1 pathway during experimental acute kidney injury. Kidney international, 73: 1240–1250, 2008.

37. van der Tuin, K, Tops, CMJ, Adank, MA, Cobben, JM, Hamdy, NAT, Jongmans, MC, Menko, FH, van Nesselrooij, BPM, Netea-Maier, RT, Oosterwijk, JC, Valk, GD, Wolffenbuttel, BHR, Hes, FJ, Morreau, H: CDC73-Related Disorders: Clinical Manifestations and Case Detection in Primary Hyperparathyroidism. J Clin Endocrinol Metab, 102: 4534–4540, 2017.

38. Inker LA, Coresh J, Levey AS, Tonelli M, Muntner P: Estimated GFR, albuminuria, and complications of chronic kidney disease. Journal of the American Society of Nephrology: JASN, 22: 2322–2331, 2011.

39. Gu L, Liang X, Wang L, Yan Y, Ni Z, Dai H, Gao J, Mou S, Wang Q, Chen X, Wang L, Qian J: Functional metabotropic glutamate receptors 1 and 5 are expressed in murine podocytes. Kidney international, 81: 458–468, 2012.

40. Collett VJ, Collingridge GL: Interactions between NMDA receptors and mGlu5 receptors expressed in HEK293 cells. BrJ Pharmacol, 142: 991–1001, 2004.

